# A FMRF-amide peptide that regulates cell non-autonomous protein homeostasis in *C. elegans*

**DOI:** 10.1101/2025.03.14.643343

**Authors:** Carrie A. Sheeler, Jacqueline Y. Lo, Daviana Menendez Escalera, Emmett Griffin-Baldwin, Karmen Mah, Adam Roszczyk, Antoine E. Roux, Cynthia Kenyon, Jennifer L. Garrison, Ashley E. Frakes

## Abstract

The coordination of protein homeostasis from the brain to periphery is essential for the health and survival of all animals. In *C. elegans*, glia serve a central role in coordinating organismal protein homeostasis and longevity via the unfolded protein response of the endoplasmic reticulum (UPR^ER^). However, the full extent of the cell non-autonomous response and the identity of the signaling molecules required remained unknown. Here, we show that glial UPR^ER^ activation induces robust transcriptomic changes in specific tissue types across the animal, particularly in pathways related to neuropeptide signaling. We performed neuropeptidomics and loss and gain-of-function genetic screens and identified a single neuropeptide, FLP-17, that is sufficient but not necessary to induce cell non-autonomous activation of the UPR^ER^. FLP-17 is sufficient to protect against chronic ER stress and age-dependent protein aggregation. We determined that FLP-17 acts through the receptor, EGL-6, to activate cell non-autonomous UPR^ER^. This work reveals a complex peptidergic signaling network initiated by glial activation of the UPR^ER^ to regulate organismal protein homeostasis.

## Introduction

All animals encounter stressors that threaten protein homeostasis, such as pathogens, insufficient or excessive nutrients, and temperature fluctuations. To sense and respond to these threats, animals have evolved cellular stress responses to restore homeostasis. The central nervous system plays a central role in coordinating the activation of cell stress responses across tissues. Activation of cell stress responses in neurons and glia induces cell non-autonomous activation of the stress response in distal peripheral tissues^1-7^. This coordination from the brain to periphery is required to coordinate appropriate physiological and behavior responses to restore homeostasis ^8,9^.

One main stress response coordinated from the brain to periphery is the unfolded protein response of the endoplasmic reticulum (UPR^ER^)^1,3-5^. The UPR^ER^ consists of three canonical branches defined by their distinct transmembrane regulator (IRE1, ATF6, and PERK). Each branch initiates specific downstream signal transduction pathways, culminating in a response that reduces synthesis of new polypeptides and increases the ER’s capacity for folding and degradation^10^. Activation of the ancestral IRE1 pathway leads to the unconventional splicing of the mRNA encoding the transcription factor X-box binding protein 1(XBP1) by the IRE1 endoribonuclease ^11,12^. Spliced XBP1 (XBP1s) regulates a range of transcriptional targets important for restoring ER protein and lipid homeostasis^13^.

During aging, there is an organism-wide loss in the ability to mount an effective UPR^ER^ which drives protein aggregation, chronic ER stress, and ultimately, tissue damage and disease susceptibility^14^. In *C. elegans*, the age-dependent decline in the ability to induce UPR^ER^ is prevented by the ectopic expression of constitutively active *xbp-1s* selectively in four astrocyte-like glial cells (*glial::xbp-1s*). *Glial::xbp-1s* induces a robust cell non-autonomous activation of the UPR^ER^ in distal cells which extends lifespan, reprograms peripheral lipid metabolism, and renders animals resistant to protein aggregation and chronic ER stress^1,15^. However, the full extent of the cell non-autonomous response and the identity of the signaling molecules required remains unknown.

Mutants deficient in neuropeptide processing and secretion suppress the cell non-autonomous activation of the UPR^ER^ and lifespan extension in *glial::xbp-1s* animals, suggesting the signal is a neuropeptide^1^. Neuropeptides are small, evolutionarily ancient neuroactive peptides that act through G Protein coupled receptors (GPCRs) to regulate many brain-controlled functions in all organisms including energy metabolism, behavior, and reproduction^16^. The *C. elegans* genome contains at least 159 predicted neuropeptide genes in three major families-insulin-related peptides (INS), neuropeptide-like proteins (NLP), and FMRF-like peptides (FLP). Together, these neuropeptide genes encode over 300 peptides, many with known or predicted GPCR binding partners^17-20^. Unlike neurotransmitters, neuropeptide secretion is not restricted to synaptic sites and can act on tissues at varying distances and timescales, making them intriguing candidates for a long-range cell non-autonomous signal^16^.

Because neuropeptide signaling can induce widespread and diverse responses in downstream tissues, we aimed to characterize the full extent of the cell non-autonomous response in *glial::xbp-1s* animals. Using whole animal single cell RNA-seq (scRNAseq), we identified robust transcriptional changes in most tissue clusters with an enrichment in genes associated with neuropeptide signaling. To identify the cell non-autonomous signal mediating these complex transcriptional changes, we performed tandem label-free quantitative mass spectrometry revealing increased levels of peptides encoded by eight genes. Loss of individual neuropeptide candidate genes failed to suppress glial-mediated UPR^ER^ activation. However, we identified one neuropeptide, FLP-17, that is sufficient to induce cell non-autonomous activation of the UPR^ER^, increase ER stress resistance, and protect against age-dependent protein aggregation. FLP-17 is secreted by two neurons (BAG neurons) and requires the receptor EGL-6 to activate intestinal UPR^ER^. Our results reveal a complex peptidergic signaling circuit involving glia and neurons to regulate organismal protein homeostasis.

## Results

### Glial *xbp-1s* induces cell non-autonomous tissue-specific transcriptional changes

To determine the tissue-specific transcriptional changes induced by glial cell non-autonomous signaling, we performed whole animal single cell RNAseq on adult glial::*xbp-1s* and control (N2) animals. For each replicate, we isolated cells from approximately 15,000 Day 1 adult animals per genotype (Figure 1A). High-level clustering revealed 20 unique cell and tissue types including neurons, socket and sheath glia, intestine, muscle, and reproductive tissues (Figure 1B). All groups demonstrated marked within-cluster separation of control and *glial::xbp-1s* cells, suggesting robust transcriptomic changes induced by *xbp-1s* expression in glia (Supplemental Figure 1A). We performed differential gene expression analysis of each tissue cluster in *glial::xbp-1s* animals compared to controls and identified differentially expressed genes (DEGs) across 15 of the 20 tissue clusters. Most tissue clusters (12/15) showed a greater number of upregulated DEGs, with only 3 tissues (germline, coelomocytes, and spermatheca) demonstrating a higher percentage of downregulated DEGs (Supplemental Figure 1B). Neurons, seam cells, and germline exhibited the most DEGs with 2863 total DEGs identified in neurons (1678 upregulated, 1185 downregulated), a total of 767 DEGs in seam cells (610 upregulated, 157 downregulated), and 730 total DEGs identified in germline (217 upregulated, 513 downregulated).

**Figure 1:**
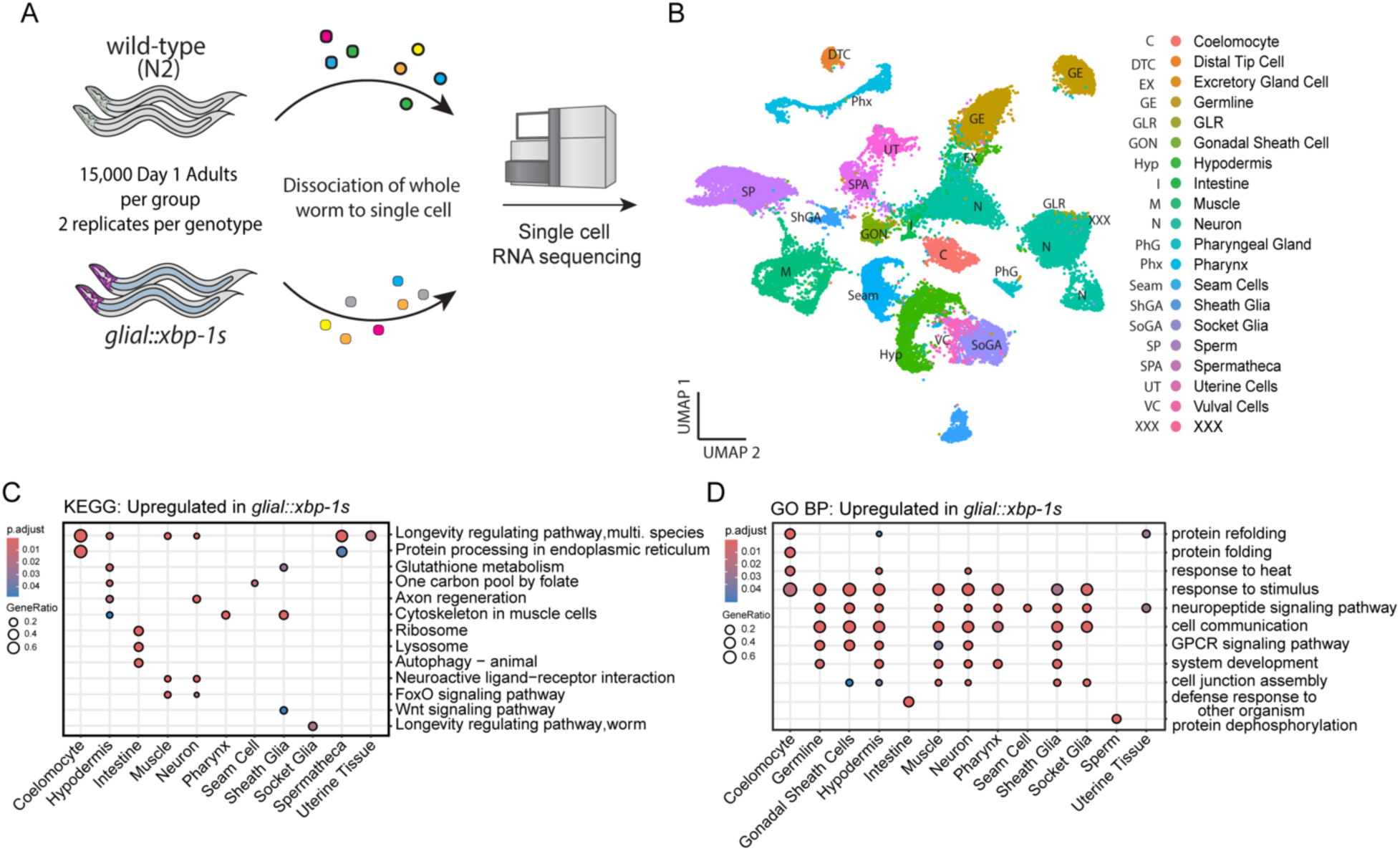
*Glial::xbp-1s* induces tissue-specific transcriptomic changes across the whole animal. (A)Whole animal single cell RNA sequencing performed on Day 1 adult *glial::xbp-1s* and wild-type (N2) animals. (B) UMAP clustering identified 20 distinct tissue and cell types which were used for differential gene expression analysis. (C) KEGG pathway enrichment demonstrated upregulation in longevity-associated pathways across multiple tissue systems and specific upregulation of autophagy and lysosome genes in the intestine of *glial::xbp-1s* animals. (D) GO analysis of biological pathways upregulated in *glial::xbp-1s* reveals that neuropeptide signaling is increased throughout the worm including in neurons, glia, germline, muscle, and hypodermis.

To identify key pathways induced by cell non-autonomous signaling by *glial::xbp-1s*, we performed KEGG and gene ontology (GO) enrichment analysis of DEGs in each tissue cluster. Based on previous whole animal bulk RNAseq and UPR^ER^ fluorescent reporter strains, we expected to observe an upregulation of UPR^ER^-dependent genes in CEPsh glia, the intestine, and pharynx of *glial::xbp-1s* animals^1^. We confirmed a trend of increased expression of XBP-1-dependent genes in the sheath glia cluster (which contains CEPsh glia), the intestine, and the pharynx, similar to the tissue-specific pattern of the UPR^ER^ reporter, *hsp-4::GFP* (Supplemental Figure 1C)^1^. However, the upregulation of some genes such as *xbp-1* were not statistically significant, likely due to technical limitations associated with the cell dissociation protocol or due to limited cell sample size for those clusters. Surprisingly, hypodermis, neurons, spermatheca, uterine tissue, and coelomocytes showed an increase in genes associated with ER function and ER stress such as protein processing in the ER (KEGG: cel04141), protein folding/refolding (GO:0006457, GO:0042026), and response to heat (GO:0009408) (Figure1C and 1D).

In addition to the UPR^ER^, *glial::xbp-1s* induces cell non-autonomous reprogramming of peripheral lipid metabolism and activation of autophagy^15^. Knockdown of genes required for autophagy suppresses lifespan extension in *glial::xbp-1s* animals^15^. We confirmed the upregulation of 6 lysosomal and proteolysis associated genes previously found to be increased in whole worm bulk RNAseq within the intestinal scRNAseq cluster (Supplemental Figure 1 D). This was further supported by KEGG enrichment analysis which revealed an increase in lysosome (KEGG: cel04142) and autophagy-associated genes (KEGG: cel04140) in the intestine (Figure 1C). Intriguingly, we also found an enrichment for defense response genes within the intestine that overlapped with previous bulk sequencing of *glial::xbp-1s* worms (Supplemental Figure 1D)^15^. These data reproduce previous findings and position the intestine as an important target tissue of glial-mediated cell non-autonomous signaling.

Previously, we demonstrated that cell non-autonomous activation of the UPR^ER^ and lifespan extension is dependent on neuropeptide signaling^1^. Therefore, we expected to observe transcriptional responses related to neuropeptide signaling by scRNA-seq. GO and KEGG enrichment analysis of DEGs showed an enrichment of neuropeptide (GO:0007218) and GPCR (GO:0007186) signaling in half of the identified tissue clusters (10 tissues out of 20 total clusters) including neurons, sheath and socket glia, muscle, and hypodermis (Figure 1C and 1D, Supplemental Figure 2A and 2B). This suggests that *xbp-1s* expression in glia activates a complex neuropeptide signaling network across many tissues.

**Figure 2.**
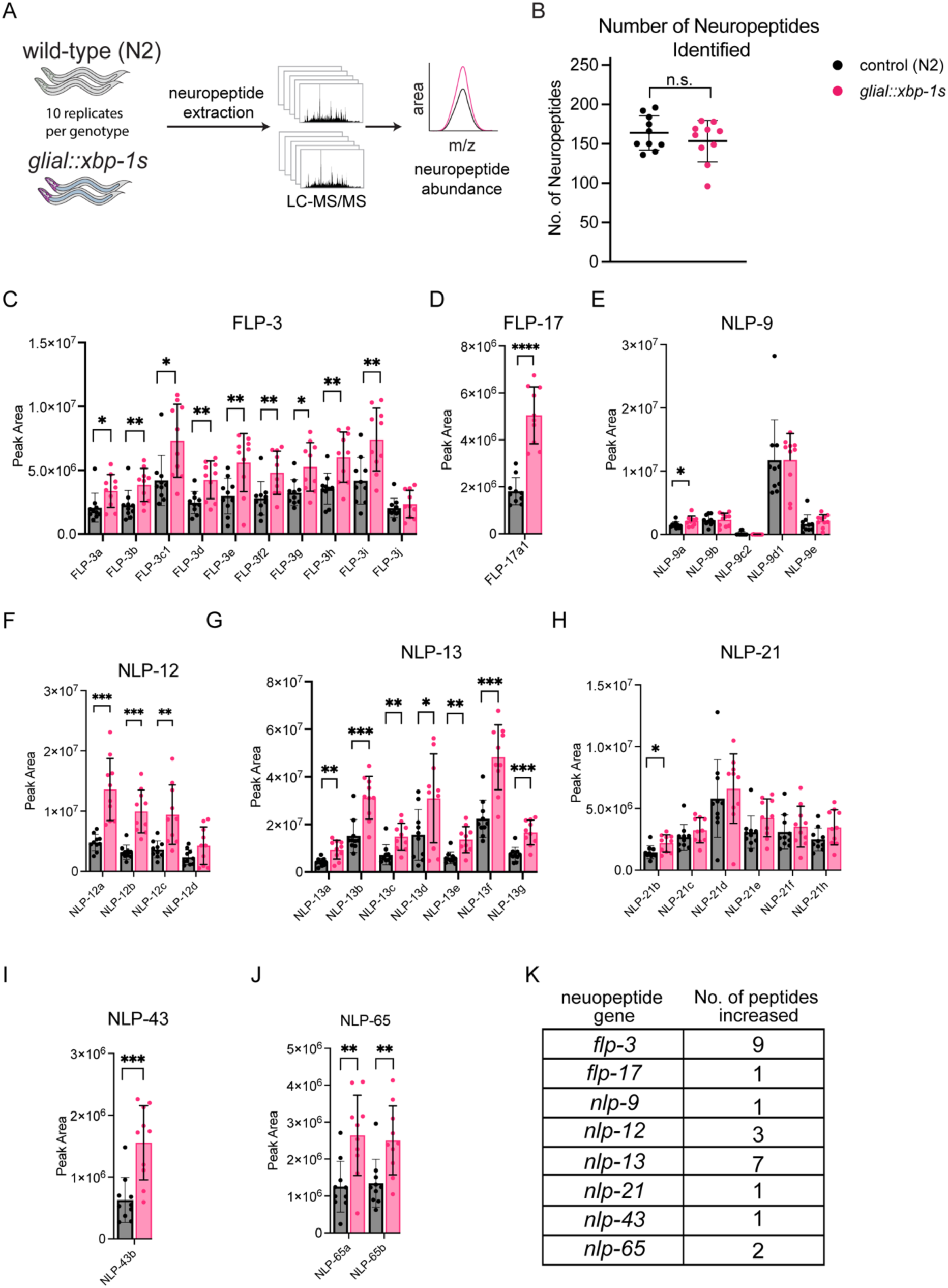
Specific neuropeptides are increased in *glial::xbp-1s* animals. (A) Schematic of neuropeptidomic analysis performed in *glial::xbp-1s* animals vs. age-matched controls (N2). resulted in a list of 11 neuropeptides dysregulated in *glial::xbp-1s*. (B) Equivalent numbers of neuropeptides were identified per genotype (each dot represents an independent biological replicate of 50,000 worms, n = 10). (C-J) Relative quantification of neuropeptides by LC-MS/MS. Peak area for each replicate is plotted and compared between two strains (n=10 biological replicates of 50,000 worms). (C) *flp-3*, (D) *flp-17*, (E) *nlp-9*, (F) *nlp-12*, (G) *nlp-13*, (H) *nlp-21*, (I) *nlp-43*, (J) *nlp-65* (* p < 0.05, ** p < 0.01, *** p < 0.001, **** p < 0.0001 student’s t-test). (K) The 25 peptides increased in *glial::xbp-1s* animals compared to controls are encoded by 8 genes.

Additionally, our data revealed unexpected transcriptional responses in tissues that we previously did not consider to be involved in cell non-autonomous signaling by *glial::xbp-1s*. For example, seven tissue clusters showed an upregulation of genes associated with longevity, including muscle, neurons, socket glia, and reproductive tissues (KEGG: cel04213 and cel04212, Figure 1C). Coelomocytes, endocytic immune-like cells in *C. elegans*, demonstrated a remarkable increase in UPR^ER^-associated pathways including protein processing in the ER (KEGG: cel04141), response to heat (GO:0009408) and protein refolding (GO:0042026). Neurons exhibited an enrichment of DEGs in FoxO signaling (KEGG: cel04068), axon regeneration (KEGG: cel04361), cell communication (GO:0007154), and a decrease of genes regulating fatty acid metabolism (KEGG: cel01212) (Figure 1C, 1D, Supplemental Figure 2A-C). This suggests that lifespan extension in *glial::xbp-1s* animals is the culmination of transcriptomic changes across multiple tissues coordinated by neuropeptide signaling.

### FLP-17 is sufficient but not necessary for cell non-autonomous activation of the UPR^ER^

To identify the neuropeptides required for cell non-autonomous activation of the UPR^ER^ and lifespan extension, we performed label-free quantitative mass spectrometry (neuropeptidomics) of *glial::xbp-1s* and control animals (Figure 2A). Neuropeptides are genetically encoded as large polypeptide precursors (pro-peptides) that are enzymatically processed by a series of enzymes to generate multiple bioactive peptides^16,20^. We detected an equivalent number of neuropeptides in *glial::xpb-1s* and control animals, confirming consistent neuropeptide isolation among the 10 biological replicates (Figure 2B). We found a total of 29 peptides significantly changed (25 increased, 4 decreased) in *glial::xbp-1s* compared to control (Figure 2C-K, Supplemental Figure 3A-D). We hypothesized that the cell non-autonomous signal driving UPR^ER^ activation would be a peptide with increased expression in *glial::xbp-1s* animals because neuropeptide deficient mutants block intestinal UPR^ER^ activation. The 25 candidate peptides increased in *glial::xbp-1s* are encoded by eight genes in the *flp* and *nlp* gene families (*flp-3*, *flp-17*, *nlp-9*, *nlp-12*, *nlp-13*, *nlp-21*, *nlp-43*, and *nlp-65).* Consistent with known limitations of this method, we were not able to identify any insulin-like peptides^17,21^.

We sought to determine if any candidate neuropeptide was sufficient to activate the UPR^ER^. To test this, we expressed each neuropeptide under its endogenous promoter (such as *flp-17p::flp-17* or *flp-17oe*) and measured UPR^ER^ activation by *hsp-4p::GFP* fluorescence (Figure 3A). We confirmed that each gene exhibited an expected expression pattern based on previously described fluorescent reporters and single cell transcriptome data (Supplementary Figure 4 and Supplementary Figure 5) ^22-24^. Interestingly, we observed only one neuropeptide, FLP-17, that was sufficient to induce cell non-autonomous activation of the UPR^ER^ in the anterior and posterior intestine by approximately 4-fold compared to controls (Figure 3B and 3C).

**Figure 3:**
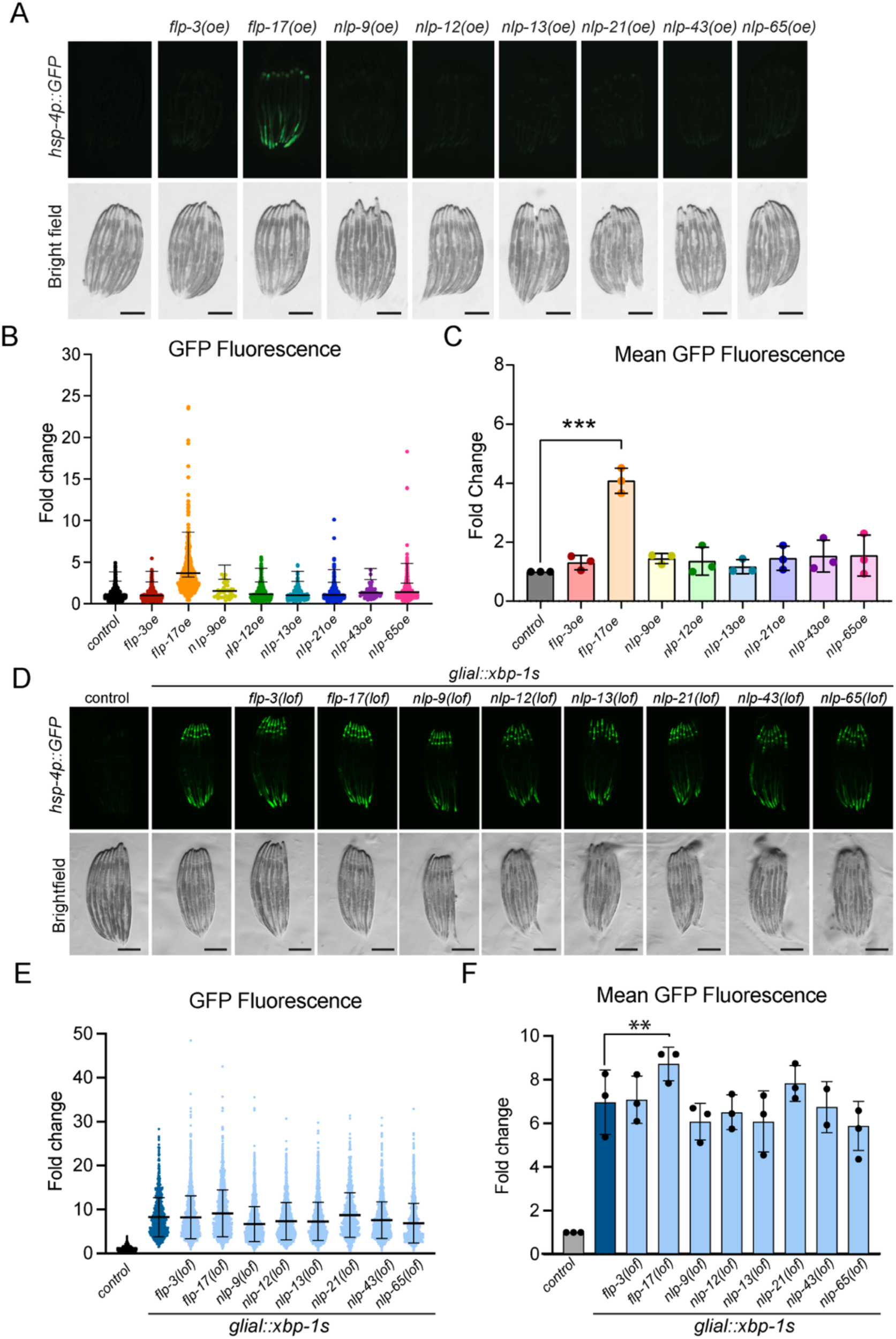
FLP-17 is sufficient to induce cell non-autonomous UPR^ER^ activation in the intestine. (A) Fluorescent micrographs of *hsp-4p::GFP* animals overexpressing each neuropeptide candidate under the endogenous promoter shows increased GFP fluorescence in the intestine of flp-17oe animals indicative of UPR^ER^ activation (scale bar = 250um). (B-C) Quantification of GFP fluorescence in neuropeptide overexpression animals by COPAS biosorter. (B) Representative experiment (dots represent individual worms). (C) Means of experimental replicates compared to *hsp-4p::GFP* control (dots = sample means from independent experiments, **** p < 0.0001, n = 3, mean ± SD, One-way ANOVA Dunnett’s Multiple Comparisons). (D) Fluorescent micrographs of *glial::xbp-1s;hsp-4p::GFP* animals harboring loss-of-function mutations in candidate neuropeptide genes (*flp-3, flp-17, nlp-9, nlp-12, nlp-21, nlp-43, and nlp-65*. (E-F) Quantification of GFP fluorescence with COPAS biosorter normalized to time of flight (length) and extinction (thickness) of animals. (E) Representative experiment (each dot represents 1 animal, error bars represent mean ± SD) and (F) means of each experimental replicate. Each dot = sample mean from one independent experiment, One-way ANOVA, Dunnett’s Multiple Comparisons, n = 2-3, ** p = 0.0041). All scale bars = 250um

To determine which candidate neuropeptides are required for cell non-autonomous activation of UPR^ER^, we introduced loss-of-function (lof) mutations for individual neuropeptide genes in animals expressing both *glial:xbp-1s* and a fluorescent UPR^ER^ reporter (*hsp-4p::GFP*) (Figure 3 D). We observed no significant changes in fluorescent intensity in *nlp-9, nlp-12, nlp-13, nlp-21, nlp-43*, or *nlp-65* when compared to *glial::xbp-1s*. Interestingly, *flp-17* mutants demonstrated a slight but significant increase in *hsp-4p::GFP* fluorescence (One way ANOVA, Dunnett’s multiple comparisons *glial::xbp-1s* vs *glial::xbp-1s;flp-17null*, p = 0.0041) (Figure 3E-F). We hypothesize that the loss of a single neuropeptide gene did not substantially suppress cell non-autonomous UPR^ER^ activation due to redundant signaling by compensatory or convergent peptidergic signaling.

### FLP-17 increases resilience to chronic ER stress and protein aggregation

Because FLP-17 induced intestinal activation of the UPR^ER^, we hypothesized that FLP-17 would extend lifespan and increase resistance to chronic ER stress. However, we observed no difference in lifespan of *flp-17oe* animals compared to controls (Figure 4A, p = 0.6912, Log-rank Mantel-Cox test). To determine if *flp-17oe* animals were more resilient to chronic ER stress, we assayed their survival when grown from adulthood on plates containing tunicamycin, an inhibitor of N-linked glycosylation that induces protein misfolding and ER stress. Interestingly, *flp-17oe* animals survived longer on tunicamycin compared to controls, demonstrating FLP-17 enhances organismal ER stress resistance (Figure 4B) (median survival 10 days compared to 8 days in control, Log-rank Mantel-Cox test p<0.0001).

**Figure 4:**
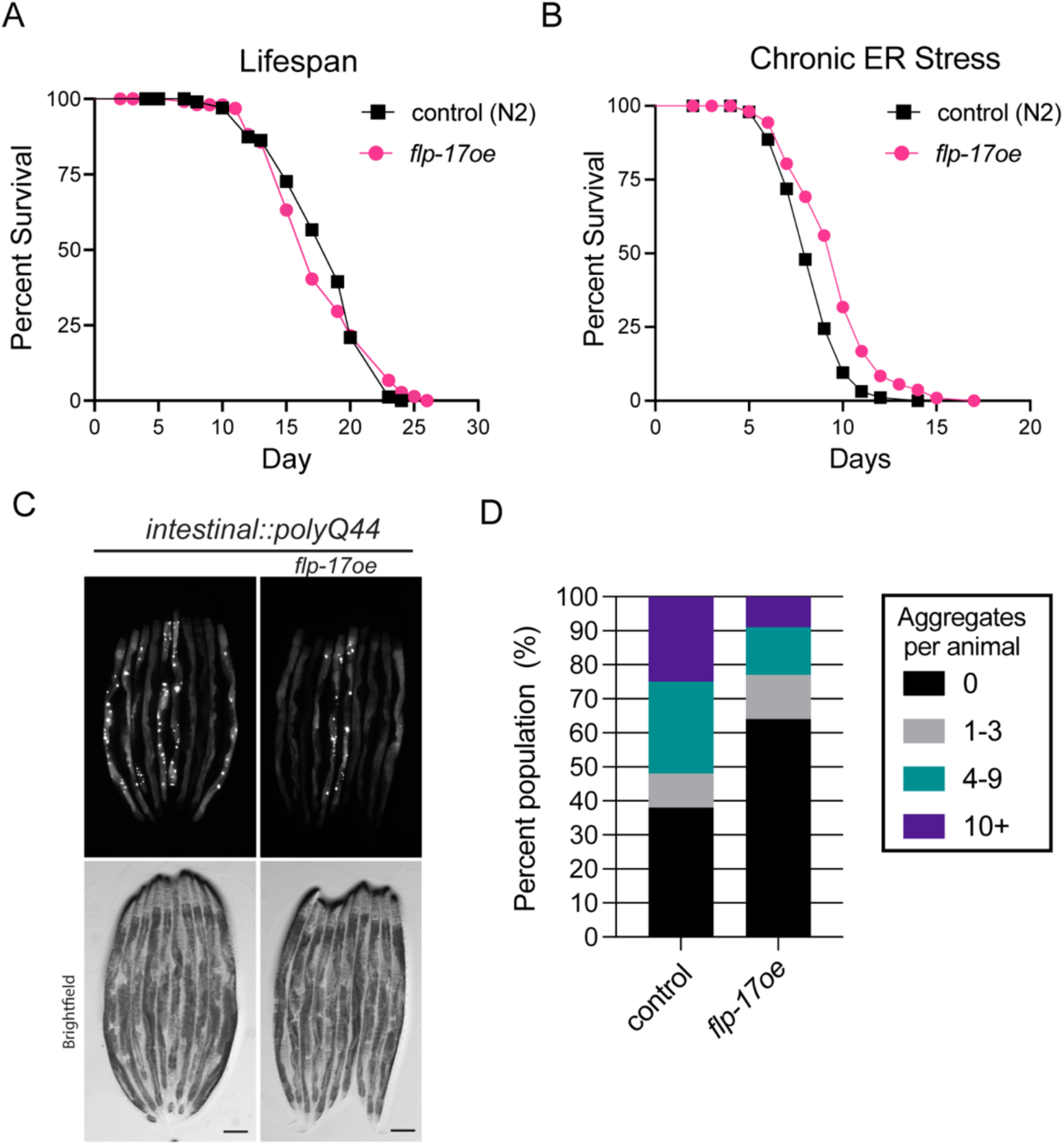
FLP-17 protects animals from chronic ER stress and protein aggregation. (A) Survival of *flp-17oe* (*flp-17p::flp-17*, pink line) vs controls (N2) (black line) (p = 0.6912, Log-rank (Mantel-Cox) test). (B) Survival of *flp-17oe* (pink line) and controls (N2) (black line) transferred to tunicamycin-containing plates at Day 1 of adulthood (p < 0.0001, Log-rank (Mantel-Cox) test). (C-D) Fluorescent micrograph (C, scale bar = 250um) and quantification (D) of polyQ44-YFP aggregation in *flp-17oe* and control animals. Number of punctae per animal quantified in Day 3 adults (n=86, controls and n=96, *flp-17oe*).

In addition to increased chronic ER stress resistance, we hypothesized that FLP-17 would confer protection against protein aggregation. Therefore, we crossed *flp-17oe* animals to a strain expressing a Huntington-like polyglutamine repeat protein (Q44) tagged with yellow fluorescent protein (YFP). These animals form intestinal protein aggregates that increase with age, visualized as YFP fluorescent puncta^25^. *Flp-17oe* reduced the number of aggregates in the intestine at day 3 of adulthood compared to controls (Figure 4C and 3D). Together, these data demonstrate that FLP-17 enhances organismal protein homeostasis via UPR^ER^ to protect animals from chronic ER stress and aging-associated protein aggregation.

### FLP-17 signals through EGL-6 to activate cell non-autonomous UPR^ER^

Next, we sought to determine which receptors FLP-17 acts through to mediate cell nonautonomous activation of the UPR^ER^. FLP-17 is secreted by two neurons (BAG neurons) and binds the G protein-coupled receptor (GPCR), EGL-6, on HSN and URX neurons to inhibit egg laying and to regulate intestinal lipid stores, respectively^26,27^. A recent large scale reverse pharmacological screen revealed FLP-17 peptides can also bind to the FRPR-8 and DMSR-1 receptors *in vitro^18^*. Therefore, we crossed *flp-17oe* animals into loss-of-function mutants for *egl-6, frpr-8*, and *dmsr-1* and measured activation of the UPR^ER^. Loss of *dmsr-1* or *frpr-8* did not suppress *flp-17*-mediated UPR^ER^ induction. However, loss of *egl-6* significantly reduced UPR^ER^ induction in *flp-17p::flp-17* animals, suggesting FLP-17 acts through EGL-6 to activate cell non-autonomous UPR^ER^ ( Figure 5A-C), unpaired t-test p = 0.0472).

**Figure 5:**
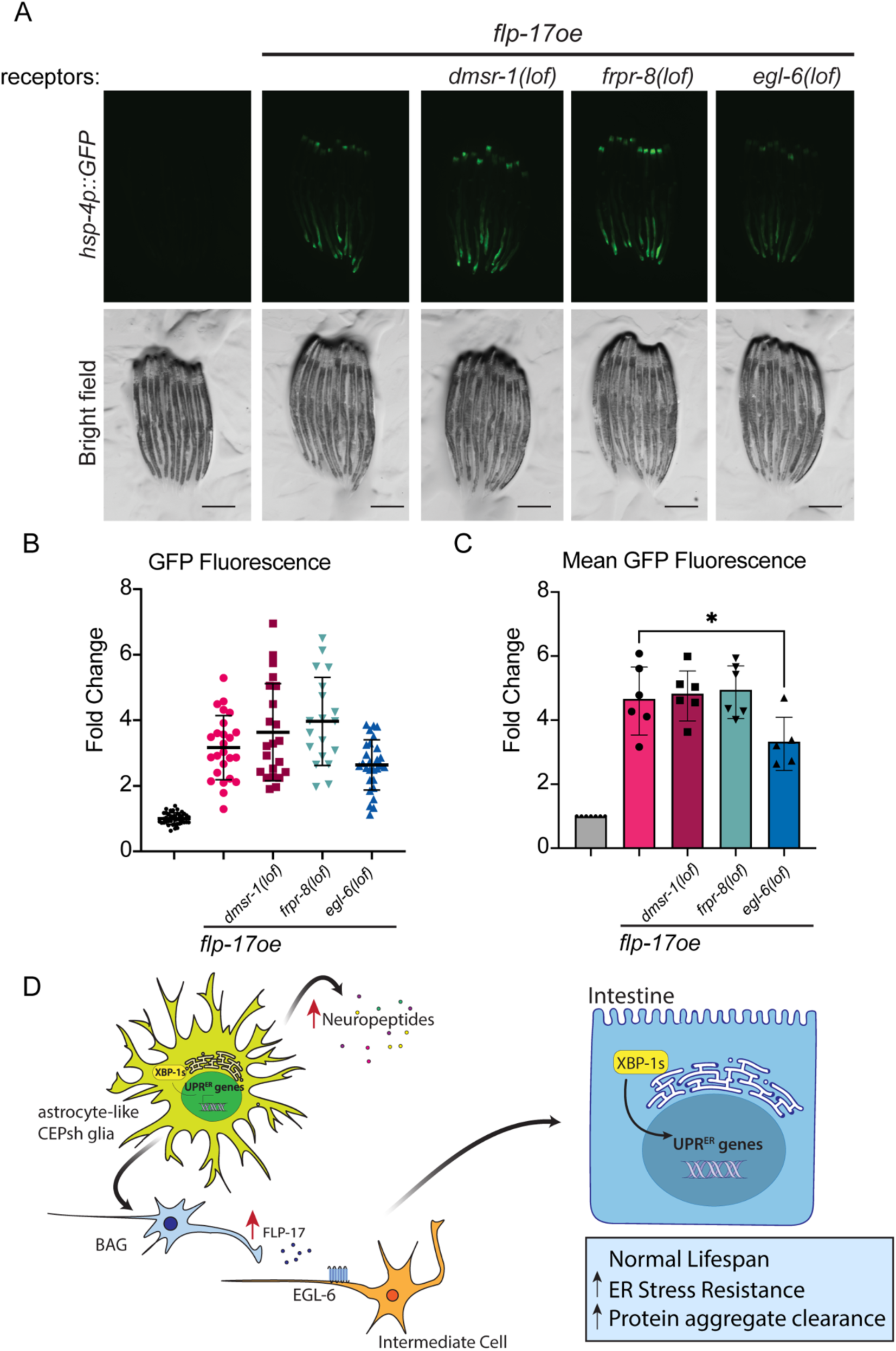
UPR^ER^ activation by FLP-17 requires the EGL-6 receptor. (A) Fluorescent micrographs of *flp-17oe;hsp-4p::GFP* animals harboring loss-of-function mutations in FLP-17 receptors: *dmsr-1, egl-6*, and *frpr-8*. (scale bar = 250um). (B) Fluorescent intensity in the posterior of adult worms was quantified using defined regions of interest (ROIs) in ImageJ and normalized to the average of *hsp-4p::GFP* controls (dots represent individual animals within one imaging experiment, n = 19-41). (C) Means of independent imaging experiments (dots represent the sample mean of an experiment, n = 5-6, * p = 0.0476, unpaired student’s t-test comparing *flp-17oe* vs *flp-17oe* with *egl-6* null). (D) Schematic of proposed model. Activation of the UPR^ER^ in CEPsh glia induces robust neuropeptide signaling. One peptide, FLP-17, is secreted by BAG neurons, binds EGL-6 on an intermediate cell which leads to intestinal UPR^ER^ activation and protection from chronic ER stress and protein aggregation.

Based on published RNA-seq and fluorescent microscopy *egl-6* is expressed in HSN motor neurons, GLR glia, and several other neurons including DVA, URX and SDQ, but has negligible to no expression in the intestine^26,28,29^. From our scRNA-seq data set, we confirmed *egl-6* expression in HSN, URX, and DVA and observed expression in other neurons including AVH, RIC, and AFD. We found no detectable expression levels of *egl-6* within GLR glia or intestine by scRNA-seq (Supplementary Figure 7A). To predict which cells FLP-17 may be acting on, we performed differential expression analysis on *egl-6*-expressing neurons in *glial::xbp-1s* animals vs controls. We observed 11 DEGs in HSN neurons, 11 in AIN, 15 in AFD, 19 in RIC, 42 in AVH, and 48 in DVA neurons (Supplementary Figure 7B-G). No differential gene expression changes were identified in GLR glia (Supplemental Figure 1B). However, this may be due to poor representation of GLR glia in the control population. Notably, AVH, RIC, and DVA neurons demonstrate increased expression of neuropeptide transcripts which may suggest altered signaling by these neurons in *glial::xbp-1s* animals. We did not observe any DEGs in the URX, AWA, or AIM neurons. Therefore, we hypothesize that within *glial::xbp-1s* animals, FLP-17 is acting through specific intermediate cell types to mediate cell non-autonomous activation of the UPR^ER^ in the intestine (Figure 4D). Collectively, these data reveal that FLP-17 is sufficient induce cell non-autonomous activation of the UPR^ER^ through the EGL-6 receptor to increase organismal protein homeostasis.

## Discussion

Our results reveal that activation of the UPR^ER^ in glial cells leads to robust tissue-specific transcriptomic changes across the animal. This coordination of UPR ^ER^ activation from the brain to peripheral tissues culminates in increased resilience to ER stress, remodeled lipid metabolism, and increased lifespan^1,15^. To identify the cell non-autonomous signal mediating these phenotypes, we performed neuropeptidomics and loss and gain-of-function genetic screens. We identified one peptide, FLP-17, that was significantly upregulated in *glial::xbp-1s* animals. Increased expression of *flp-17* was sufficient to induce cell non-autonomous UPR^ER^, protect against chronic ER stress and protein aggregation.

FLP-17 belongs to an evolutionarily conserved class, FMRF-amide/RF-amide neuropeptides, that play important roles in energy balance and reproduction across phyla^30,31^*. In C. elegans,* FLP-17 is secreted from a pair of sensory neurons (BAG) in response to low oxygen and high carbon dioxide which can be caused by unfavorable food conditions or pathogens^32,33^. FLP-17 then acts through specific neurons to inhibit egg laying and initiate an aversion behavior until the animal has reached more favorable conditions^26,32^. Interestingly, unfavorable food conditions and pathogens also threaten organismal protein homeostasis^34,35^. Therefore, we speculate that FLP-17 evolved to simultaneously protect the animal from proteotoxic stress while facilitating a behavioral program to help the animal navigate to more favorable conditions.

Although FLP-17 was sufficient to activate the UPR^ER^, it was not necessary for cell non-autonomous activation of the UPR^ER^ by *glial::xbp-1s.* We speculate that the inability for the *flp-17* null mutant to suppress cell non-autonomous activation of the UPR^ER^ is due to neuropeptide redundancy or genetic compensation. For example, *glial::xbp-1s* might induce multiple peptides that are functionally redundant in activating the UPR^ER^. Besides FLP-17, no other candidate peptide from the neuropeptidomics screen was sufficient to induce the UPR^ER^, suggesting the candidate peptides are not sufficient to compensate for loss of *flp-17*. However, we cannot rule out compensation by a peptide not included in our list of candidates, such as insulin-like peptides. It is also possible that loss of a single neuropeptide gene could induce genetic compensation of an alternate neuropeptide^36^. Genetic compensation of neuropeptide genes was observed in zebrafish to ensure that normal gonad development and reproductive capacity continue in the absence of gonadotropin-releasing hormone and kisspeptin^37^.

We determined that FLP-17 acts through EGL-6 to induce cell nonautonomous activation of the UPR^ER^ in the intestine. However, transcriptional reporters and single cell RNAseq show negligible to no expression of *egl-6* in the intestine. This suggests FLP-17 is acting through an intermediate cell type to activate the intestinal UPR^ER^ ^26,28^. Previous work demonstrated that FLP-17 can regulate intestinal lipid metabolism by acting through URX neurons. FLP-17 binds EGL-6 on URX neurons, reducing URX activity and suppressing intestinal fat loss^27^. Interestingly, we did not detect any differential expressed genes in URX neurons. Future work is needed to elucidate how FLP-17/EGL-6 signaling leads UPR^ER^ activation in the intestine.

Overall, this study provides insight into how cellular stress resistance can be coordinated across an entire animal. We identified a specific FMRF-amide peptide, FLP-17, that is sufficient to increase organismal resilience to proteotoxic stress. Our findings reveal a new avenue to promote healthy aging and prevent or delay chronic age-onset disease.

## Methods

### Worm Maintenance

*C. elegans* were maintained at 15°C or 20°C on standard nematode growth medium (NGM) agar plates seeded with *Escherichia coli* OP50-1. All experimentation was performed at 20°C on animals fed with OP50-1. All animals were age-matched by bleach syncing and assayed at Day 1 of adulthood unless otherwise specified. The N2 Bristol strain was used as a reference for wild type.

### Worm Strains

Three neuropeptide null strains were created through CRISPR-Cas9 driven excision of the entire gene sequence of *nlp-9*, *nlp-13*, and *nlp-65* as previously described^38,39^. In brief, gRNAs predicted by TargetScan were designed to target PAM sites identified at the 5’ and 3’ ends of the target sequence. gRNAs and Cas9 were ordered from IDT, sequences listed in Table 2. Successful injections were determined by co-CRISPR with dpy-10 which was selected against when isolating neuropeptide mutants. Homozygous neuropeptide deletions were confirmed by PCR-based genotyping.

**Table 1.**
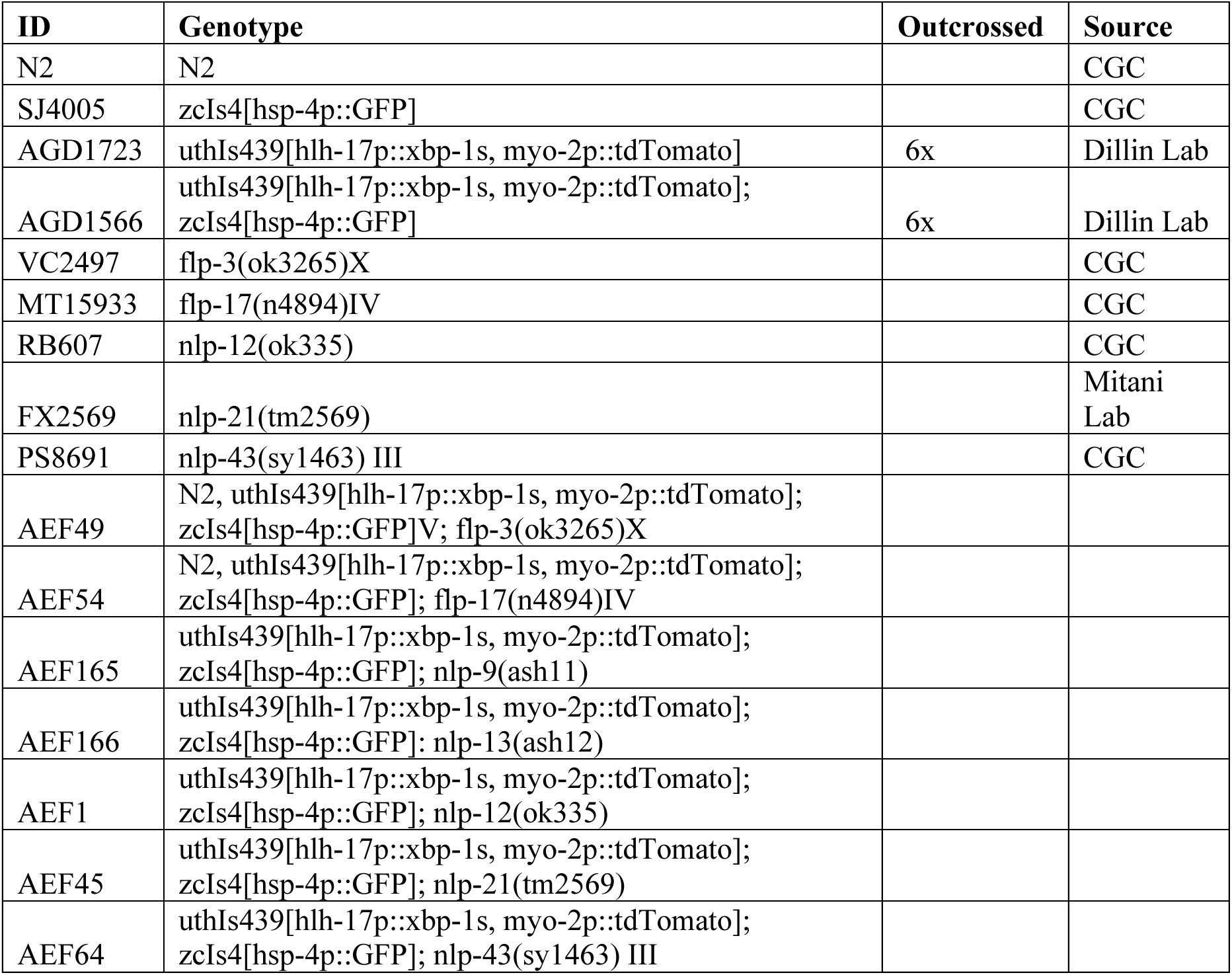
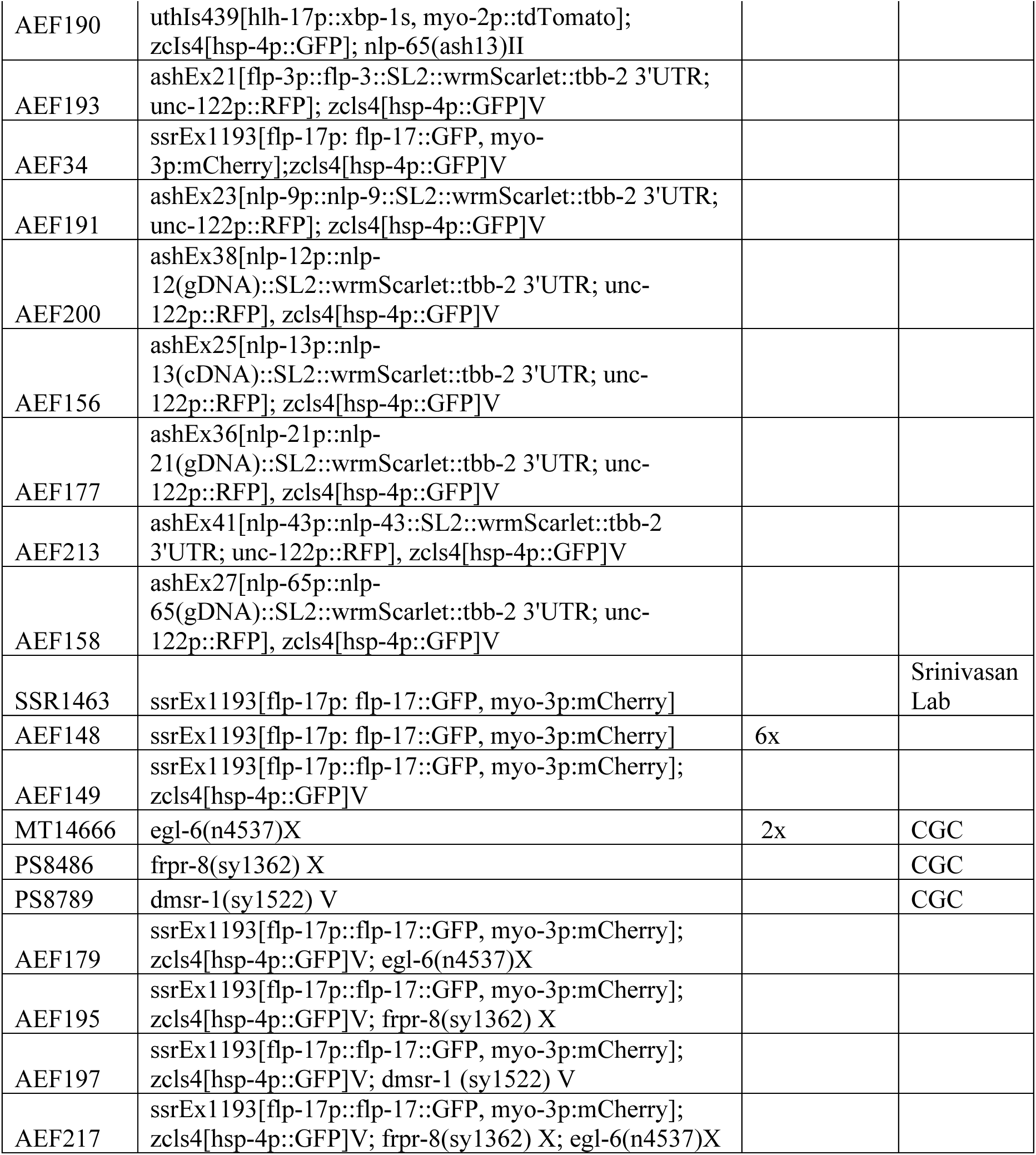
Worm Strains.

**Table 2.**
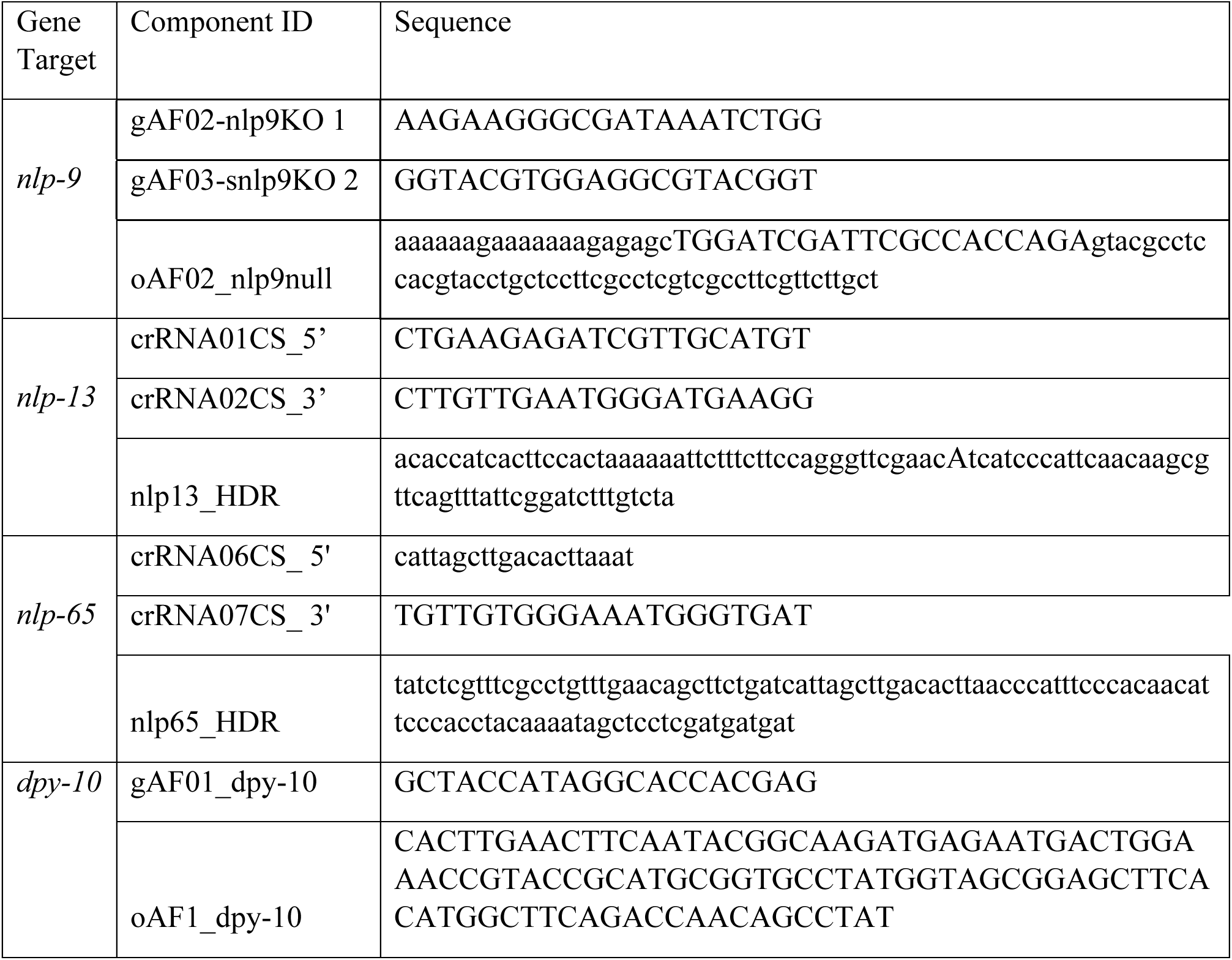
gRNAs and HDR oligos.

Extrachromosomal arrays for *flp-3*, *nlp-9*, *nlp-12*, *nlp-13*, *nlp-21*, *nlp-43*, and *nlp-65* were designed to drive the expression of each gene behind their endogenous promoter. Promoter regions were based on previous literature or the 5’ region from neuropeptide start codon until the next upstream gene (Table 3).

**Table 3.**
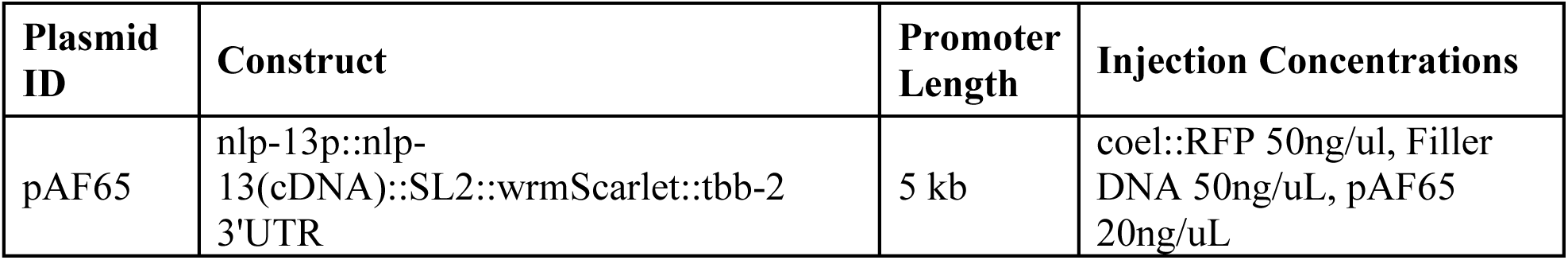
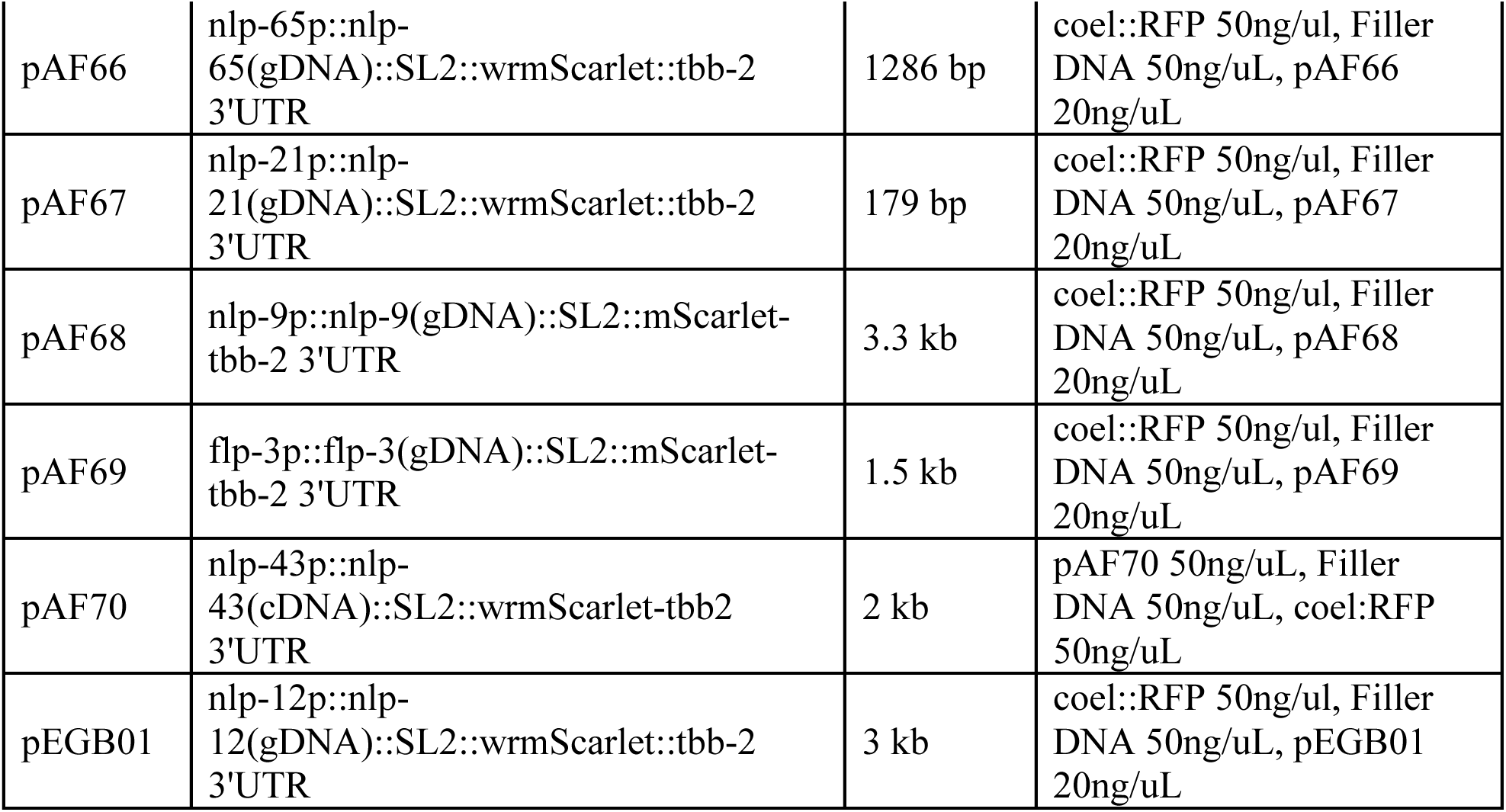

### Plasmids

To create the pAF65 plasmid for nlp-13oe, the *nlp-13* promoter sequence was amplified from genomic DNA (gDNA), combined with cDNA for the *nlp-13* gene (gAF08) and gene block for SL2::wrmScarlet (gbAF09), and cloned via Gibson assembly into the pAF61 backbone. Similarly, for nlp-21oe, the 3kb promoter sequence was amplified from gDNA and cloned into the pAF61 backbone with gene blocks *nlp-21* cDNA (gbAF07) and SL2::wrmScarlet (gbAF10). The nlp-65oe encoding array (pAF66) was assembled from a 2.3kb gDNA fragment containing both the 1286bp promoter and the 1014bp *nlp-65* gene, cDNA for SL2::wrmScarlet (gbAF11), and inserted into the pAF61 backbone. *nlp-9* (pAF68), *flp-3* (pAF69), and *nlp-12* (pEGB01) were created by amplifying gDNA containing the combined promoter and gene sequence for each gene and cloning it into the backbone of pAF67 using Gibson assembly. For pAF70, nlp-43p::nlp-43(cDNA) was subcloned into the pAF65 backbone by Azenta Life Sciences. Whole plasmids were sequence verified, then injected into SJ4005 (plasmids and injection mix, Table 3).

### Single Cell Sequencing: Cell sorting and single cell RNA sequencing

N2 and *glial::xbp-1s* animals were age-synced with bleach and harvested at Day 1 of adulthood. Samples were prepared as previously described^40^. Briefly, cells were dissociated by a chemical treatment with SDS and DTT of approximately 5 minutes to disrupt the cuticle, followed by room temperature incubation in a pronase solution in L15 Leibovitz medium. Cells were dissociated by passage through a 21G syringe until cells were dissociated. After dissociation, the cell suspension was fixed in methanol and centrifuged 7.5 minutes at 500 rcf 4°C. The cells were gently re-hydrated with 2 mL egg buffer BSA with 2U/µL RNase inhibitor. DAPI was added at 3 µM and incubated for 10 minutes on ice. The cells were filtered through a 40 µM strainer and sorted with BDFACS Aria Fusion with the 100µm nozzle running at a frequency of 31.0 kHz. To reduce the amount of germline, only the 2N DNA population was gated and sorted into 1mL PBS BSA 2x RNAse inhibitor 4U/µl. The sorting solution was transferred to a 5 mL tube and centrifuged 7.5 minutes at 500 rcf 4°C, the pellet was resuspended in PBS BSA 10 mg/ml with 2U/µl RNAse inhibitor. 16,000 cells were loaded into a 10X Genomics Chromium chip to aim for 10,000 cells per channel. Final libraries were processed on an Illumina cBot and HiSeq4000 sequencer.

### Single Cell Sequencing: Analysis

Four samples each containing ∼15,000 worms, two representing the N2 background and two from the *glial::xbp-1s* strain, were sequenced with paired end reads. Initial processing and alignment of reads was performed on the NIH HPC Biowulf cluster (https://hpc.nih.gov). In brief, FASTQ files for each sample were aligned to *C. elegans* reference transcriptome WBcel235(ensembl109) using 10x Genomics CellRanger v7.1.0. Subsequent analysis was performed in RStudio (R v4.3.2). Quality of resulting cells was determined as follows. Likely dead or dying cells were identified by > 4% expression of mitochondrial DNA genes including *nduo-6*, *atp-6*, *nduo-2*, *ctb-1*, *ctc-3*, *nduo-4*, *ctc-1*, *ctc-2*, *nduo-3*, and *nduo-5* and removed from analysis. DoubletFinder v2.0 was then used to identify and remove cells which represented likely multiples^41^. Finally, ambient RNA contamination was eliminated using DecontX^42^. Single cell data was scaled and normalized using Seurat v4.3.0^43^. Because DoubletFinder is less able to identify doublets from the same cell type, cells with abnormally high gene counts or features were removed from analysis as potential doublets. Similarly, any cells remaining in the dataset with fewer than 500 feature counts were removed as potential empty wells. The Louvain algorithm was used to perform an initial clustering of the dataset into 22 tissue-level groups.

Cluster identification was performed using the top 10 genes which defined each “tissue”. Top genes were compared against established, published datasets and the Enrichment Analysis tool on the *C. elegans* database Wormbase to confirm the identity of given clusters^28,40,44-46^. Notably, several of these clusters had the same marker genes in common, such as clusters 1 and 11 which exhibited characteristic expression of neuronal marker genes and were subsequently named with the same identity. Each tissue-level group was further separated into single cell types using the Leiden algorithm^47^. This was most effective for neurons and glia, which have highly distinct transcriptional profiles and which have been well characterized by previous sequencing publications^28,46^. Other tissue types, such as hypodermis and seam cells, were too transcriptionally homogenous to separate into subpopulations of cells during subclustering. As a potential note of interest, most reproductive tissues were found to cluster in a similar area of the UMAP. However, vulval cells were always found to cluster with socket glia. Single cell types which clustered with other “tissues” such as vulval cells with socket glia or GLR glia with neurons were marked as distinct populations on the final tissue level dataset leaving a final count of 20 distinct tissue level clusters.

To determine the effect of *glial::xbp-1s* on transcriptional profiles, differential gene expression analysis was run using DESeq2 with the ashr package to reduce false discovery rates^48,49^. Gene ontology and KEGG enrichment analyses on resulting lists of differentially expressed genes were performed using clusterProfiler v4.13.4^50,51^. Top pathways of interest were manually selected for graphical representation (Figure 1C and 1D) with full lists of pathways provided (Supplementary Figure 2A, 2B, and 2C)

### Neuropeptide isolation

Biochemical isolation of endogenous neuropeptides from C. elegans was performed as previously described^21,52^. Strains were grown from L1 to adulthood in liquid culture, according to standard protocol (WormBook), with OP50 in the liquid culture at OD600 1.68, at 20°C, shaking at 200 rpm. Worms were collected at day 2 of adulthood for mass spectrometry analysis. Ten replicates of 50,000-60,000 worms were used per strain. Day 2 adult hermaphrodite worms were collected from culture, washed 3-5 times with M9 to remove bacteria. All liquid was removed before flash-freezing samples in liquid nitrogen and storing at - 80°C. Frozen worm pellets were dounce homogenized in 10 ml of 90:9:1 methanol/water/acetic acid, followed by probe sonication (15 seconds on, 45 seconds off; 4 cycles total at 80% amplitude with 3 mm ultrasonic probe tip, Cole-Parmer cat. no. SI-04712-12) after transferring lysates to a 15 ml conical tube. Worm lysates were then centrifuged at max speed at 4°C (Eppendorf 5810R centrifuge), and the supernatant was saved. Methanol was removed using a speedvac (reducing volume from 10 ml to 1 ml). The 1 ml of sample was transferred to low-binding 1.5 ml tubes (Thermo Fisher cat. no. 3451) then centrifuged again at max speed at 4°C (Eppendorf 5430R centrifuge). The supernatant was transferred to a new glass culture tube and then delipidated with equal volume hexane (Sigma cat. no. 139386) in the glass culture tube, twice, recovering the aqueous phase each time and combining. The aqueous fraction containing the neuropeptides was passed through a size-exclusion filter (GE Healthcare sephadex PD MidiTrap G-10 cat. no. 28918011) to remove molecules < 700 Da. Lastly, samples were desalted using a C18 spin column (Thermo Fisher cat. no. 89873) and stored in the stated elution buffer at 4°C before mass spectrometry analysis. Unless otherwise stated, low-binding 1.5 ml tubes were used throughout the protocol after lysis to mitigate loss of sample.

### LC-MS/MS

To detect neuropeptides, samples were run on a Waters nanoACQUITY coupled online to a Thermo Scientific Q Exactive Plus mass spectrometer. The EASY-Spray analytical column was used: EASY-Spray column, 15 cm x 75 µm ID, PepMap C18, 3 µm; Thermo Scientific ES800. The gradient used was: flow rate of 300 nL/min, 3-8% acetonitrile with 0.1% formic acid over 5 minutes, 8-20% acetonitrile with 0.1% formic acid over 50 minutes, 20-36% acetonitrile with 0.1% formic acid over 10 minutes. Mass spectrometer settings: data-dependent (dynamic exclusion settings at 25 s) Top10 method (charge selection excluding unassigned, 1); HCD (Higher-energy collisional dissociation); full MS scan resolution of 70,000, with a maximum injection time of 100ms and scan range of 400 to 1600 m/z; MS/MS scan resolution of 17,500, with a maximum injection time of 120 ms.

### Mass spectrometry data analysis

PEAKS Studio X and the Q module (Bioinformatics Solutions, Inc.) was used to anlayze LC-MS/MS. The data was searched against a database containing: all *C. elegans* proteins (Wormbase), E. coli proteins (SwissProt), and common protein contaminants (CRAPome89). Parent mass error tolerance was set at 20 ppm and fragment mass error tolerance was set at 0.03 Da. Enzyme was set at “none” and digest mode to “unspecific.” Variable modifications included: amidation (-0.98), enzymatic glycine removal leaving an amidated C-terminus (-58.01), half of a disulfide bridge (-1.01), oxidation on M (+15.99), phosphorylation on STY (+79.97), pyro-glu from E (-18.01), and pyro-glu from Q (-17.03). Label-free quantification analysis was performed using the Q module of the PEAKS software, using TIC intensity selected as normalization. Only peptides that were detected and differentially changed in all ten replicates of each strain were considered in the candidate list.

Biochemical isolation of endogenous neuropeptides from age-synced day 2 *C. elegans* was performed as previously described ^21^. In brief, LC/MS of neuropeptide samples was performed on a Waters nanoACQUITY coupled online to a Thermo Scientific Q Exactive Plus mass spectrometer using an EASY-Spray analytical column, 15 cm x 75 μm ID, PepMap C18, 3 μm; Thermo Scientific ES800. LC-MS/MS data were analyzed using PEAKS Studio X and the Q module (Bioinformatics Solutions, Inc.). The data was searched against a database containing all C. elegans proteins (Wormbase), E. coli proteins (SwissProt), and common protein contaminants (CRAPome). Variable modifications included: amidation (-0.98), enzymatic glycine removal leaving an amidated C-terminus (-58.01), half of a disulfide bridge (-1.01), oxidation on M (+15.99), phosphorylation on STY (+79.97), pyro-glu from E (-18.01), and pyro-glu from Q (- 17.03). Label-free quantification analysis was performed using the Q module of the PEAKS software, using TIC intensity selected as normalization.

### Fluorescent Microscopy and Image Quantification

Representative fluorescent microscopy was performed at day 1 of adulthood on worms that were anesthetized with 0.3% w/v sodium azide (NaN_3_) using a Leica M205FCA fluorescent stereo microscope. 10 worms per group are included for each image. All worms compared in an analysis were aligned together on the same NGM plate. Each experiment was performed a minimum of 3 times.

To quantify regional GFP fluorescence, 50-100 array+ worms were collected with M9, anesthetized and imaged on a 100mm NGM agar plate using a Leica M205FCA fluorescent stereo microscope. Integrated density of GFP fluorescent in the posterior half of the worm was measured using ImageJ. Regions of interest (ROI) were defined by brightfield as the area from the vulva to the tip of the tail. ROI areas were consistent across sampled populations. Fold change of GFP fluorescence and area for each sample population was determined relative to SJ4005 controls. This analysis was performed 5-6 times per strain. The mean fluorescence from each experimental replicate was used to determine statistical significance through an unpaired student’s t-test.

Imaging to capture cell-specific neuropeptide expression was performed on anesthetized day 1 adult animals using a Zeiss LSM 780 confocal microscope.

### Large Particle Flow Cytometry (COPAS)

A complex object parameter analysis sorter (COPAS) Biosort (Union Biometrica, no. 350-5000-000) was used to measure GFP fluorescence of individual worms for quantification. Total integrated GFP fluorescence was normalized to time of flight (length) and extinction (thickness) of each individual animal. Each data point represents an individual animal measurement. Each COPAS experiment was performed a minimum of 2 times. Data was analyzed using a one-way ANOVA followed by a Tukey post-hoc analysis.

### Lifespan Assay

Lifespan analyses were performed on NGM plates at 20°C on OP50-1 *E. coli*. Animals were age-synchronized by bleaching, plated on bacteria and grown from hatch to adulthood at 20°C. Day 1 adults were moved to 12 plates containing 10 animals each. Adult worms were manually moved away from progeny onto fresh plates every day or every other day during reproductive stage of adulthood. Animals were scored every 1-2 days for death or censored due to bagging, vulval protrusion, or crawling off the plate.

Lifespans were performed blind and run a minimum of 3 times by at least 2 independent researchers with representative data presented. Prism 10 was used to for statistical analyses, p-values were calculated using long-rank (Mantel-Cox) method.

### ER Stress Resistance Assay

At day 1 of adulthood, age-synced animals were transferred to NGM plates containing either 25ug/mL of tunicamycin (Sigma T7765). Animal survival was scored daily. All assays were performed 3 independent times. Prism 10 was used for statistical analyses. p values were calculated using long-rank (Mantel-Cox) method.

### Protein Aggregate Formation

Age-synced animals were manually moved at day 1 and day 2 of adulthood to new plates to keep them separate from progeny. Fluorescent microscopy was performed at day 3 of adulthood of a minimum of 100 worms per strain. The number of YFP+ punctae was counted and recorded for each individual worm. Blinding was performed at day 1 and maintained until quantification was complete. The assay was repeated 3 times independently by two researchers.

## Acknowledgements

We thank the Srinivasan laboratory and Caenorhabditis Genetics Center (CGC) for several strains used in this study. We thank Dr. Raz Bar-Ziv for assistance during scRNA-seq cell dissociation and Han Yuan from David Kelley’s lab at Calico for initial insights and assistance with the scRNAseq data. Further information and requests for reagents may be directed to A.F.

## Author Contributions

CS designed and performed experiments, analyzed data, generated *C. elegans* strains, analyzed scRNA-seq data, and wrote the manuscript. JL performed neuropeptidomics and data analysis. DME, EGB, KM, AR, assisted in generating strains and performed experiments. AER performed scRNA-seq cell isolation. CK provided resources for scRNAseq. JLG provided resources for neuropeptidomics and provided intellectual contributions. AEF designed and performed experiments, analyzed data, generated *C. elegans* strains, performed scRNAseq cell isolation, and wrote manuscript.

