## Supplemental Figures for "A FMRF-amide peptide that regulates cell non-autonomous protein homeostasis in *C. elegans*"

A

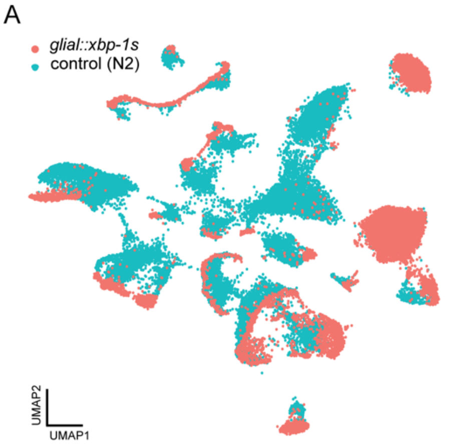


B

| | Differentially Expressed Genes | | | Cells Assayed | |
|---|---|---|---|---|---|
| **Tissue Identity** | Total | Up | Down | *glial::xbp-1s* | control |
| Coelomocyte | 86 | 23 | 63 | 220 | 1131 |
| Distal Tip Cell | 0 | 0 | 0 | 40 | 405 |
| Excretory Gland Cell | 0 | 0 | 0 | 11 | 1 |
| Germline | 730 | 217 | 513 | 1923 | 2957 |
| GLR | 0 | 0 | 0 | 139 | 3 |
| Gonadal Sheath Cell | 105 | 100 | 5 | 81 | 560 |
| Hypodermis | 718 | 534 | 184 | 632 | 1414 |
| Intestine | 118 | 94 | 24 | 84 | 305 |
| Muscle | 688 | 523 | 165 | 842 | 2316 |
| Neuron | 2863 | 1678 | 1185 | 6455 | 4049 |
| Pharyngeal Gland | 0 | 0 | 0 | 82 | 145 |
| Pharynx | 426 | 368 | 58 | 903 | 555 |
| Seam Cell | 767 | 610 | 157 | 280 | 1223 |
| Sheath Glia | 345 | 323 | 22 | 760 | 684 |
| Socket Glia | 284 | 236 | 48 | 1113 | 606 |
| Sperm | 416 | 347 | 69 | 296 | 2708 |
| Spermatheca | 107 | 42 | 65 | 124 | 810 |
| Uterine Tissue | 100 | 70 | 30 | 255 | 671 |
| Vulval Cells | 131 | 115 | 16 | 123 | 237 |
| XXX | 0 | 0 | 0 | 45 | 0 |

DEG : q < 0.05 & LogFC > absval(0.5)

C D

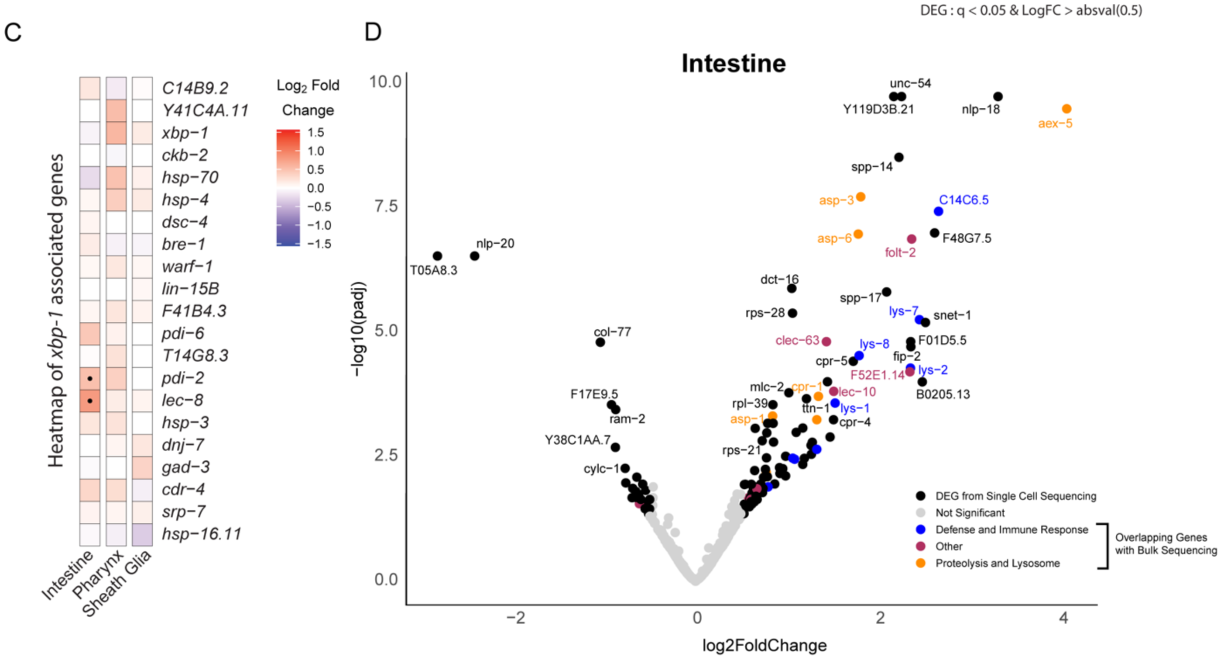


**Supplementary Figure 1: Single cell RNA sequencing in *glial::xbp-1s* animals.**
(A) Tissue level UMAP clustering of scRNA-seq data colored to highlight cells from *glial::xbp-1s* animals (orange) and controls (N2) (teal). (B) Table demonstrating differential gene expression (q < 0.05, LogFC > absval(0.5)) and number of cells assayed per condition within tissue-level clusters. (C) Heatmap of expression for known *xbp-1*-regulated genes in intestine, pharynx, and sheath glia (redder hue represents increased LogFC, dots represent genes with a q < 0.05). (D) Volcano plot of differential gene expression in the intestine. Black dots are DEGs identified in scRNA-seq, maroon, orange and blue dots are DEGs genes also identified in bulk whole animal RNAseq of *glial::xbp-1s* vs N2 (Metcalf et al. 2023), grey dots are not significant).

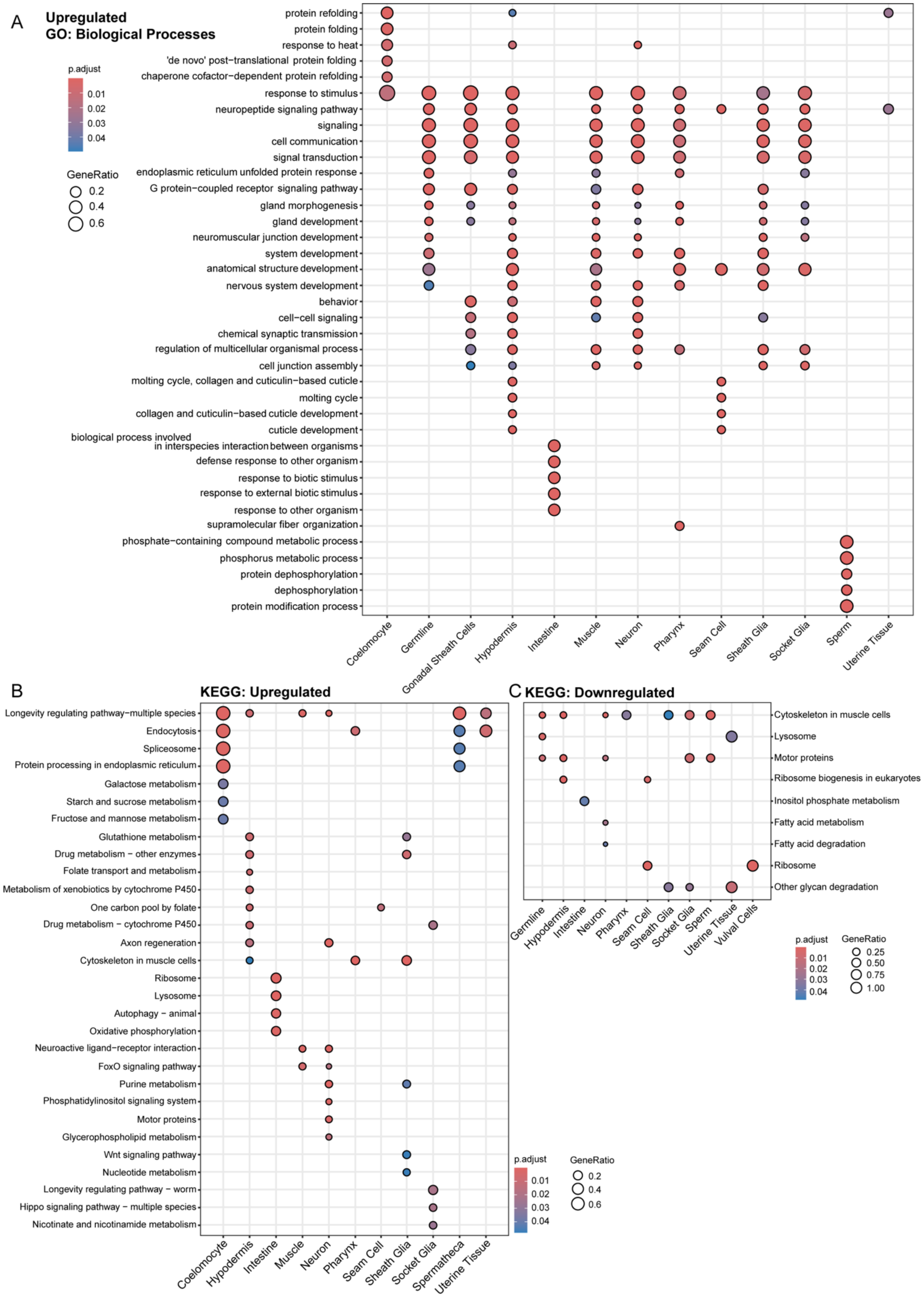


**Supplementary Figure 2: Gene Ontology and KEGG enrichment in *glial::xbp-1s*.**

(A) All top GO:BP pathways upregulated within *glial::xbp-1s* compared to control. Full graphs of KEGG enrichment analysis of upregulated (B) and downregulated (C) pathways in tissue level clusters.

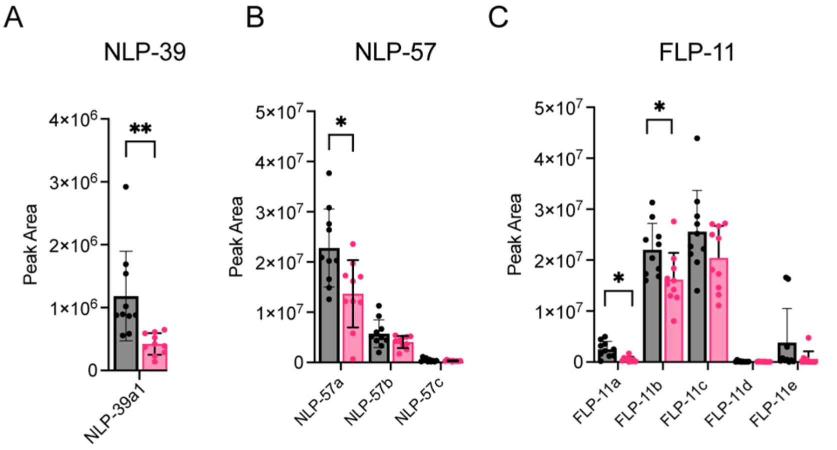


D

| Neuropeptide gene | No. of peptides decreased |
|---|---|
| *nlp-39* | 1 |
| *nlp-57* | 1 |
| *flp-11* | 2 |

**Supplementary Figure 3: Specific neuropeptides are decreased in *glial::xbp-1s* animals.**
(A-C) Relative quantification of neuropeptides by LC-MS/MS. Peak area for each replicate is plotted and compared between two strains (n=10 biological replicates of 50,000 worms). (D) The four peptides decreased in *glial::xbp-1s* animals compared to controls are encoded by 3 genes. (* $p < 0.05$, ** $p < 0.01$, student's t-test)

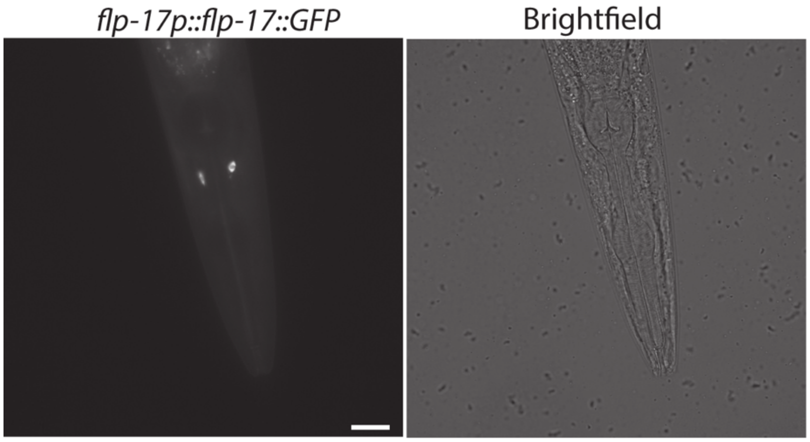


**Supplementary Figure 4: *flp-17* is expressed in BAG neurons.**
A) Fluorescent micrograph of *flp-17p::flp-17::GFP* animals confirms *flp-17* is expressed exclusively in BAG neurons as previously described[26]. Scale bar = 20um.

A

*flp-3*

| Cell type | Counts |
|---|---|
| IL1 | 1357.306 |
| VC | 847.373 |
| Hypodermis | 203.504 |
| Muscle | 96.185 |

B

*nlp-9*

| Cell type | Counts |
|---|---|
| VB | 521.548 |
| SMD | 221.663 |
| SIA | 198.657 |
| Hypodermis | 146.822 |

C

*nlp-12*

| Cell type | Counts |
|---|---|
| DVA | 3430.459 |
| Gonadal sheath cell | 872.119 |
| Hypodermis | 754.687 |
| Germline | 312.736 |

D

*nlp-13*

| Cell type | Counts |
|---|---|
| PDA | 1083.968 |
| VC | 691.569 |
| PVW | 427.027 |
| VB | 311.140 |

E

*nlp-21*

| Cell type | Counts |
|---|---|
| VB | 1352.638 |
| AVE | 561.664 |
| AVA | 550.112 |
| Hypodermis | 350.580 |

F

*nlp-43*

| Cell type | Counts |
|---|---|
| DVC | 1155.771 |
| PVW | 216.068 |
| IL neurons | 169.512 |
| Hypodermis | 80.179 |

G

*nlp-65*

| Cell type | Counts |
|---|---|
| Pharyngeal gland | 159.806 |
| SAA | 149.837 |
| Germline | 41.890 |
| AVM/PVM | 24.847 |

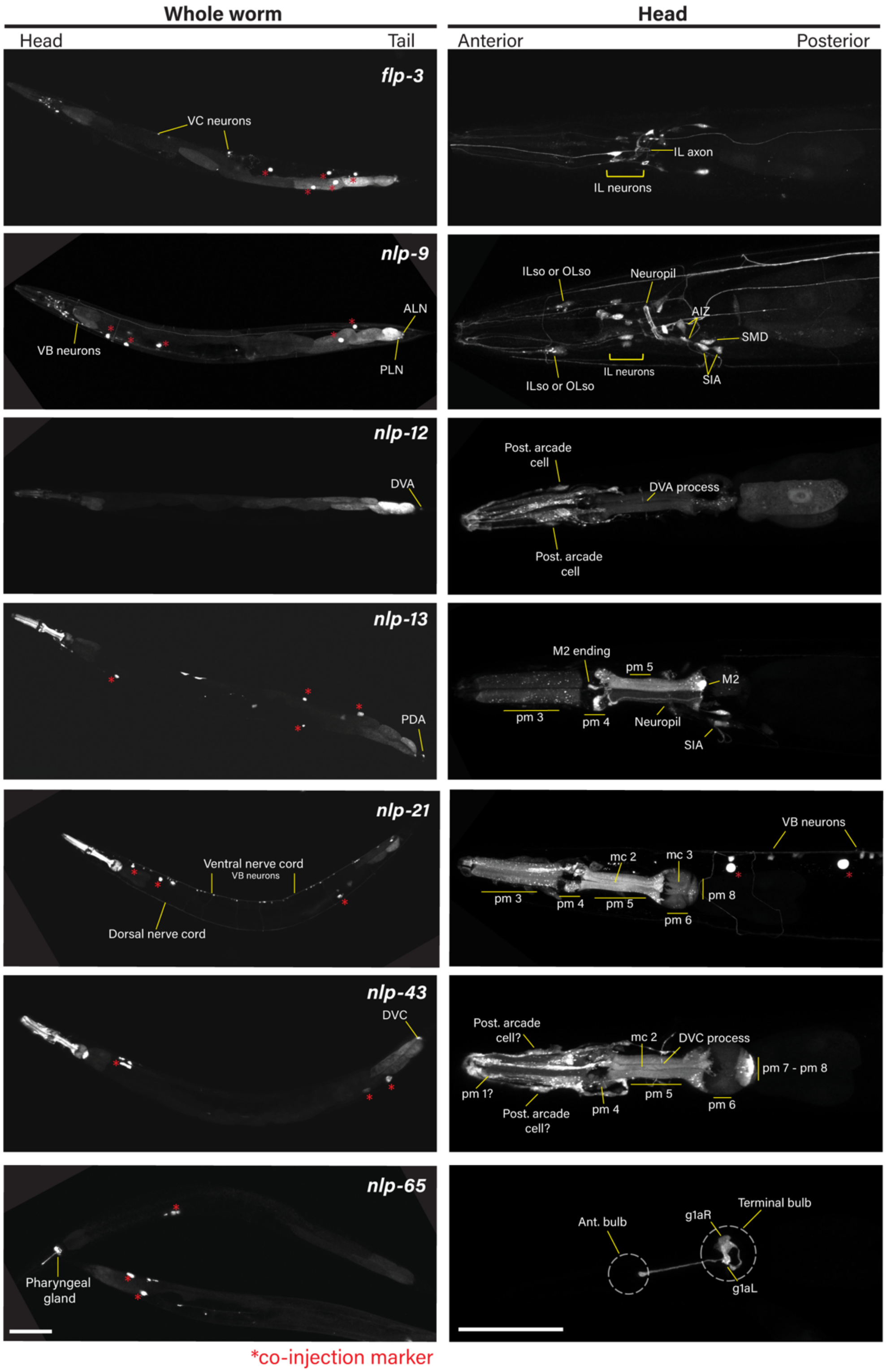


**Supplementary Figure 5: Validation of neuropeptide expression patterns.**
(A-G) Top 4 predicted sources of specific neuropeptides from scRNA-seq data reported for each strain (counts= mean counts for each gene in the given cell or tissue type) and correlating images of wrmScarlet in the whole worm (left) and head (right) of (A) *flp-3oe,* (B) *nlp-9*oe, (C) *nlp-12oe*, (D) *nlp-13oe*, (E) *nlp-21oe*, (F) *nlp-43oe*, and (G) *nlp-65oe*. Red asterisks indicate coelomocytes

expressing RFP as a co-injection marker. Scale bar = 100um for whole worm images, scale bar = 50um for head images.

A

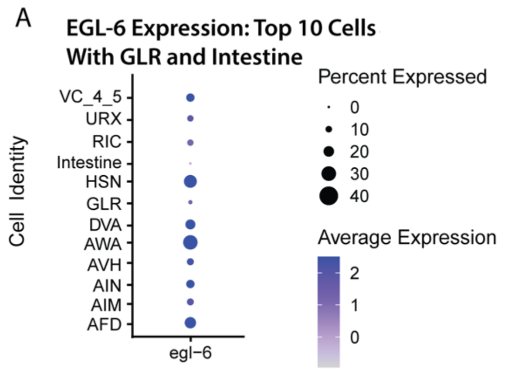


B **Differential Gene Expression: Top 10 EGL-6 Cells**

| | DEGs | | | Cells Assayed | |
|---|---|---|---|---|---|
| **Cell Identity** | Total | Up | Down | *glial::xbp-1s* | control |
| VC 4 and 5 | 2 | 1 | 1 | 27 | 56 |
| URX | 0 | 0 | 0 | 37 | 15 |
| RIC | 19 | 17 | 2 | 83 | 10 |
| HSN | 11 | 1 | 10 | 58 | 129 |
| DVA | 48 | 21 | 27 | 34 | 101 |
| AWA | 0 | 0 | 0 | 41 | 20 |
| AVH | 42 | 23 | 19 | 114 | 37 |
| AIN | 11 | 10 | 1 | 53 | 9 |
| AIM | 0 | 0 | 0 | 48 | 6 |
| AFD | 15 | 12 | 3 | 68 | 37 |

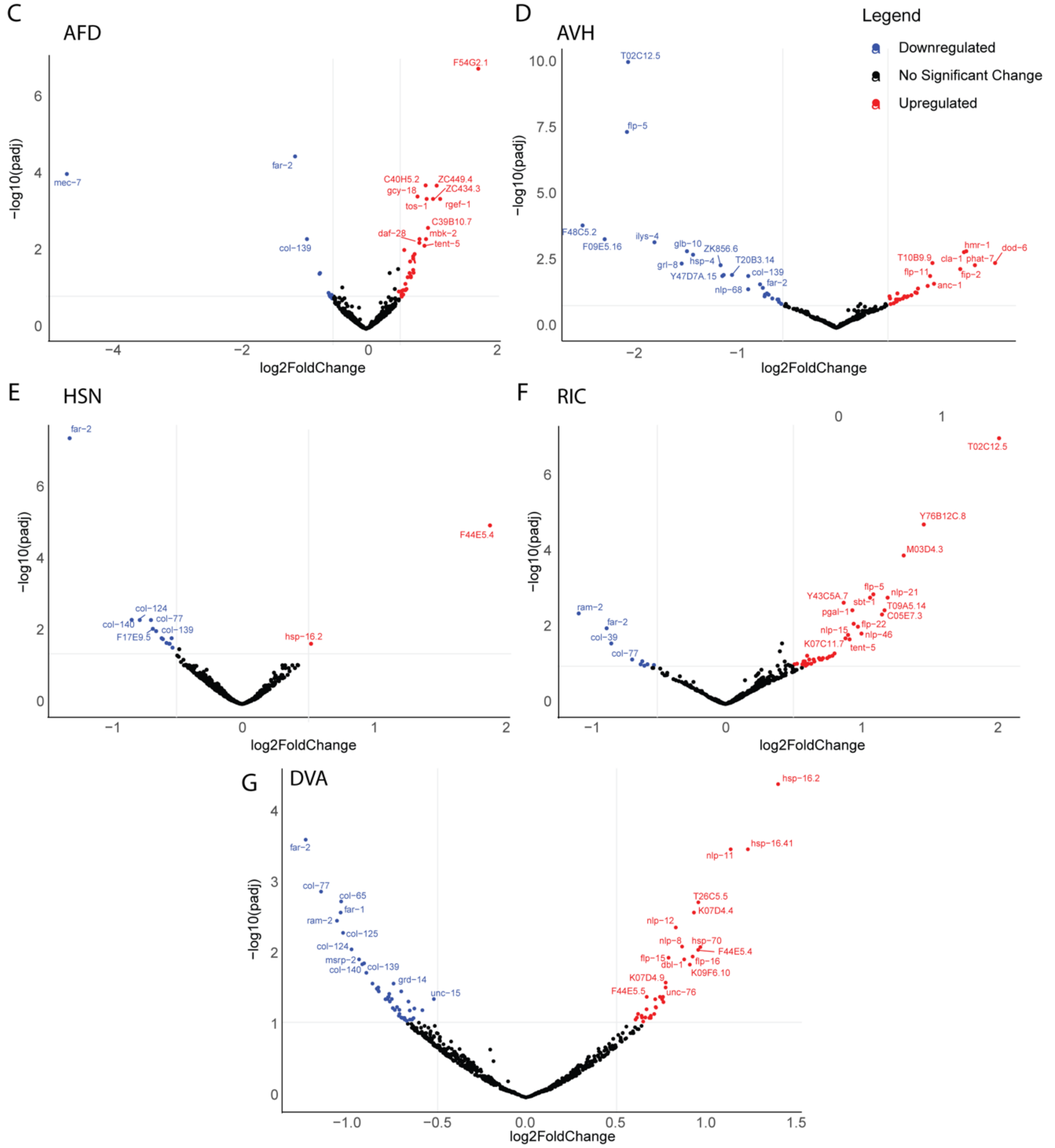


**Supplementary Figure 6: Expression of *egl-6* and cell-specific differential gene expression in top *egl-6* expressing neurons**. (A) Dot plot of GLR glia, intestine, and the top 10 neurons expressing transcripts for *egl-6* from the single cell sequencing analysis. (B) Differential gene expression analysis of the top 10 predicted neuronal expressers of *egl-6* in *glial::xbp-1s* compared to N2, controls (q < 0.05, logFC > absval(0.5)). (C-F) Volcano plots show DEGs in five neurons that likely express *egl-6* including (C) AFD, (D) AVH, (E) HSN, (F) RIC, and (G) DVA neurons (red denotes genes with q < 0.05 and LogFC > 0.5, blue denotes genes with q < 0.05 and LogFC < -0.5, black denotes no significant gene expression change. Labels are provided where legibility permits for genes that fit the q < 0.05 cutoff).
